# Structurally Constrained Effective Brain Connectivity

**DOI:** 10.1101/310938

**Authors:** Alessandro Crimi, Luca Dodero, Fabio Sambataro, Vittorio Murino, Diego Sona

## Abstract

The relationship between structure and function is of interest in many research fields involving the study of complex biological processes. In neuroscience in particular, the fusion of structural and functional data can help understanding the underlying principles of the operational networks in the brain. To address this issue, this paper proposes a constrained autoregressive model leading to a representation of “effective” connectivity that can be used to better understand how the structure modulates the function. Or simply, it can be used to find novel biomarkers characterizing groups of subjects. In practice, an initial structural connectivity representation is re-weighted to explain the functional co-activations. This is obtained by minimizing the reconstruction error of an autoregressive model constrained by the structural connectivity prior. The model has been designed to also include indirect connections, allowing to split direct and indirect components in the functional connectivity, and it can be used with raw and deconvoluted BOLD signal.

The derived representation of dependencies was compared to the well known dynamic causal model, giving results closer to known ground-truth. Further evaluation of the proposed effective network was performed on two typical tasks. In a first experiment the direct functional dependencies were tested on a community detection problem, where the brain was partitioned using the effective networks across multiple subjects. In a second experiment the model was validated in a case-control task, which aimed at differentiating healthy subjects from individuals with autism spectrum disorder. Results showed that using effective connectivity leads to clusters better describing the functional interactions in the community detection task, while maintaining the original structural organization, and obtaining a better discrimination in the case-control classification task.

**Highlights:**

- A method to combine structural and functional connectivity by using autoregressive model is proposed.
- The autoregressive model is constrained by structural connectivity defining coefficients for Granger causality.
- The usefulness of the generated effective connections is tested on simulations, ground-truth default mode network experiments, a classification and clustering task.
- The method can be used for direct and indirect connections, and with raw and deconvoluted BOLD signal.

## 1. Introduction

Investigation of how structure influences function is present in several fields. For example, how the function of a protein can be predicted by its sequence and structure is a common issue in proteomics (Lee et al., 2007; Watson et al., 2005). Network representation can aid this task by bridging diverse types of data across different domains (Oh et al., 2014). Indeed, networks are present at many scales from cell type-specific metabolic or regulatory pathways within neurons to the interactions between cortical areas, which are the focus of this paper. The human brain is a complex network characterized by spatially interconnected regions that can be activated during specific tasks or at rest. As a consequence, integration of segregated regions (communities) is emerging as the most likely structural organization explaining the complexity of brain function (see e.g. Hermundstad et al. (2013)). More specifically, *functional connectivity* refers to covarying activity between spatially segregated brain regions measured in time series data. *Structural connectivity* refers to the physical tracts connecting brain regions generally estimated in-vivo by diffusion weighted images. *Effective connectivity*, which combines structural and functional information, refers to the influence that one neural system exerts over another, either at neuronal or brain region level (Friston, 2011; Hahn et al., 2019). In this context, we propose a model that re-defines (sparsifies) structural connectivity based on functional information, to subsequently give a representation of effective connectivity. In this way, effective connections are the connections which are used to biophysical transfer activities between brain areas.

Indeed, deeper understanding of the relationship between the functional activity in different brain regions and the structural network highlighted by using tractography can convey useful information about brain underlying principles. While in Fukushima et al. (2018) and Hermundstad et al. (2013) it has been shown that there is a significant overlap between neuroanatomical connections and correlations of brain functional signals, it is yet to be fully understood how the whole-brain network interacts during specific tasks or at rest, explaining all structural and functional aspects especially in terms of causality. A predictive framework based on multiple sparse linear regression was used to predict functional series from structural data (Deligianni et al., 2013), and a platform called the “virtual brain” was designed to simulate brain activity in injured and healthy subjects (Jirsa et al., 2010). With the aim of highlighting the relationships between function and structural connectivity, a different approach relies on a Bayesian framework to estimate the functional connectivity using a structural graph as a prior (Hinne et al., 2014). However, results generating functional connectivity from structural information have been so far challenging, due to the fact that certain high correlations appear between brain regions not directly linked by structural connections. To overcome this limitation approaches based on causal graphs (Rajapakse and Zhou, 2007; Chicharro and Panzeri, 2014) and on propagators and neural field theory (Robin-son, 2012) were proposed.

Even more restricting works have criticized the view of effective connectivity as biophysical transfer of activity between brain areas, since functional activities might arise decoupled by structure and biophysical transfer and arising for reasons yet unknown (Preti and Van De Ville, 2019; Medaglia et al., 2018; Tyszka et al., 2011). Currently, there are still ongoing controversies on the definition of causal interaction between brain regions due to possible confounding properties each model can involve (Reid et al., 2019). Despite the criticisms of being mostly measures of temporal correlations (Bielczyk et al., 2018; Etkin, 2018), among the methods expressively based on causal interactions, the most popular are the dynamic causal model (DCM) (Friston, 2011) and the Granger causality (Granger, 1969). DCM uses an explicit model of regional neural dynamics to capture changes in regional activations and inter-regional coupling in response to stimulus or task demands. Statistical inference is used to estimate parameters related to directed influences between neuronal elements. While this is a powerful method to study effective connectivity, its main limitation is the combinatorial complexity on the number of modeled regions and connections, which limits its applicability to only few regions. Despite attemts to generalize to brain-wide have been proposed (Razi et al., 2017).

Although not directly, Granger causality also quantifies the causal influence and the flow of information between regions. Despite the slow dynamics and the regional variability of the hemodynamic response, making Granger causality a controversial method for the analysis of functional magnetic resonance images (fMRIs), it has been used to identify the dynamics of Blood-Oxygen-Level Dependent (BOLD) signal flow between brain regions (Goebel et al., 2003; Liao et al., 2011). Notwithstanding, spurious influences might be obtained due to the fact that the hemodynamic activity and the underlying neuronal activity have different temporal resolutions, resulting in different propagation times and delays (Rangaprakash et al., 2018). Fortunately, a proposed solution is to perform a deconvolution to remove the hemodynamic actitivities from the BOLD signal before performing causal analysis (Wu et al., 2013). Recent works have criticized both DCM and Granger causality being predictors of events identifying temporal correlation and not true biological causalities (Bielczyk et al., 2018; Etkin, 2018). Nevertheless, DCM and Granger causality are pragmatic and well defined measures of causal influence, and have delivered many insights in neuroscience and physiology (Friston, 2011; Goebel et al., 2003; Deshpande et al., 2008). Despite the criticisms, they are still the most used models of causal inference.

On the other hand, looking at the structural connectivity, the result of any tractography algorithm might be imprecise, introducing false positive connections (Chen et al., 2015). Refining the fiber tracking by keeping only the fibers that are supposed to be used during tasks or resting-state might be a way to reduce false positives. The method we propose aims at estimating the coefficients of a Granger causality taking into considerations functional and structural data. This is achieved by filtering the structural connectivity by using a multivariate autoregressive (MAR) model constrained in a novel manner: by preserving and re-weighting the structural connections that are required to parsimoniously reproduce the measured BOLD dynamics.

A MAR model is a random process specifying a linear dependence of variables from their previous values and from a stochastic term. Thank to this, Granger causality does not suffer an excessive computational complexity. Nonetheless, in case of a large number of regions involved in the analysis, the model is affected by cancellation issues and high sensitivity to noise (Stokes and Purdon, 2017). Moreover, the computed coefficients do not necessarily correspond to existing structural connections. This makes it cumbersome to perform a whole brain analysis involving many brain regions. Furthermore, Granger causality approach has not yet used structural connectivity in a multivariate manner.

Taking inspiration from the aforementioned limitations, this paper presents an extension of the MAR model introducing a constrained MAR (CMAR), which uses the structural connectivity as a prior to bound the search space during parameter fitting. This fusion of structural connectivity and functional timeseries aims at representing an effective brain connectivity, addressing a whole brain analysis thanks to the sparse representation of the connectivity matrix. In its simple form the constrained model identifies putative direct structural connections implicitly using functional signal. To include indirect connections as shown in Figure 1, the model can be expanded through neural field theory inspired propagators (Robinson, 2012) also based on tractography.

**Figure 1:**
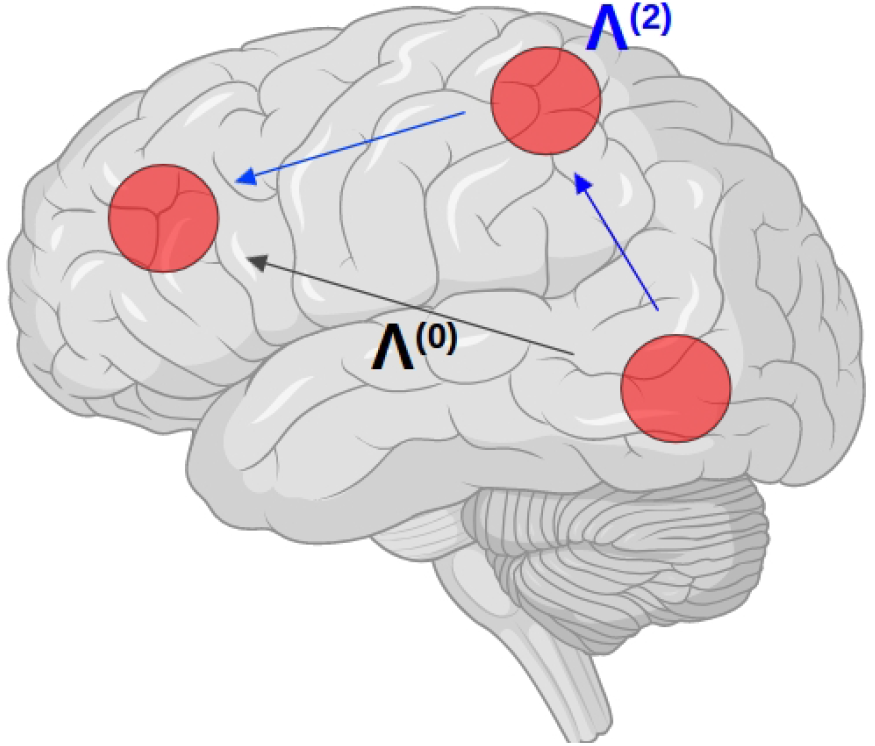
Simplistic representation of the possible propagation path from one region to another (created by using Biorender.io). Λ^(0)^ represents a direct connection, while Λ^(2)^ depicts a 2-step indirect connection.

## 2. Overview

In the Method section we first describe the used datasets, followed by the preprocessing steps and finally the technical details. This paves the way to the Results section where experiments on simulations, default mode network (DMN), classification of autism spectrum disorder (ASD) and typically developing (TD) subjects and effective community detection are then reported. Some useful background for those experiments is reported in the following paragraphs.

### 2.1. Default Mode Network

Among brain networks DMN, which is a set of brain regions more active during rest rather than a goal directed task, has emerged for its possible role in allocating attention, self-referential processing, and memory (Demertzi et al.; Greicius et al., 2003). Moreover, some of those DMN regions can belong to different large scale intrinsic networks as shown in whole-brain studies on resting-state (Yeo et al., 2014). After a simulation experiment, DMN regions are used as a proof of concept for the method.

### 2.2. Autism Spectrum Disorder

Autism is a developmental disorder characterized by repetitive, restricted behavior and deficits in communication and social interactions (Fishman et al., 2015). The neurobiology of ASD is still unclear, and the discrimination from TD subjects using neuroimaging is still difficult to perform (Yahata et al., 2016). Nevertheless, connectome-based classifiers for ASD and TD individuals have been recently introduced also characterizing local and global graph metrics of structural and functional networks (Yahata et al., 2016; Rudie et al., 2013). We applied our method on ASD dataset to validate the discriminant power of the obtained representations.

### 2.3. Community detection

Clustering group of nodes within brain networks is relevant since it can show meaningful behavior of sub-graphs across multiple samples. where those subgraphs are often called modules, clusters or communities. This is a characteristic that exists in other several real world systems including social, biological, and political systems (Fortunato, 2010). In neuroscience the related question is on how neural units cluster into densely interconnected groups that provide coordinated activities such as perception, action, and adaptive behaviors (Meunier et al., 2010). Moreover, the principle of modularity characterizes the fundamental organization of human brain functional connectivity during learning (Bassett et al., 2011), and sparsely inter-connected modules allow faster adaptation of the system in response to changing environmental conditions (Meunier et al., 2010). Therefore, in this context of parsimony, our proposed approach can lead to the identification of sparser community which can relate better functional and structural information.

## 3. Methods and Data

The aim of the proposed method is to combine structural brain connectivity and functional brain activity. To reach this goal we resort to a multivariate autoregressive model properly modified in order to allow for the estimation of the temporal brain activation biased by the structural connectivity. In all our experiments we used the Power atlas defining 264 putative functional regions of interest (ROIs) (Power et al., 2011). More specifically we used the version with non-overlapping ROIs (see below). A functional atlas has been chosen given the need to map functional series. The experiments have been conducted on publicly available datasets described below for which the ethical approval has already been granted.

### 3.1. Nathan Kline Institute-Rockland dataset

To estimate the connectome communities and analyze the DMN, a large dataset obtained by the 1000 connectomes initiative was used. This comprised 200 right-handed healthy subjects from the Nathan Kline Institute-Rockland dataset (Nooner et al., 2012), and publicly available at the url http://fcon1000.projects.nitrc.org/. For each subject, resting state fMRI (rs-fMRI), diffusion weighted imaging (DWI) and T1 were acquired by using a Siemens (Munich, Germany) Magnetom scanner and co-registered. rs-FMRI data were acquired with the eyes open, using a 3 Tesla scanner, with TR/TE times as 1.4s/30ms, flip angle 65°, and isotropic voxel size of 2 mm, for a total scan duration of 10 minutes. DWI volumes were acquired with a 1.5 Tesla scanner by using 35 gradient directions and TR/TE 2.4s/85ms, flip angle 90^°^, and isotropic voxel-size of 2.5 mm. The T1 weighted MRI data were acquired with the same 1.5 Tesla scanner, using as TR/TE times 1.1s/4.38ms, flip angle 15°, and isotropic voxel-size of 1 mm.

### 3.2. Autism Brain Imaging Data Exchange II dataset

To perform the case-control classification task, the experiments were performed on the San Diego State University cohort of the ABIDE-II dataset publicly available (Fishman et al., 2015). This cohort was chosen among the others of the ABIDE-II dataset as it has been aquired at a resolution high enough to perform tractography (Di Martino et al., 2017). This final dataset included 26 ASD and 21 TD subjects. For each subject, rs-fMRI, DWI and T1-weighted were acquired and co-registered. Imaging data were acquired on a GE (Milwaukee, WI) 3T MR750 scanner. For a detailed description of the experimental protocols refer to Fishman et al. (2015). Briefly, rs-fMRI volumes were acquired using a single-shot gradient-recalled, echo-planar pulse sequence, in one 6:10-minute eyes-open scan consisting of 185 whole brain volumes at TR/TE = 2s/30ms, flip angle 90°, and isotropic 3.4 mm voxel-size. The DWI volumes were acquired with a dual spin echo excitation using TR/TE = 8.5s/84.9ms, flip angle 90°, and 1.88 × 1.88 × 2*mm*^3^ voxel-size. Total diffusion-weighted scan time was about 9 minutes. T1-weighted inversion recovery spoiled gradient echo sequence were acquired at TR/TE = 8.1s/3,172ms, flip angle 8°, and isotropic 1 mm voxel-size.

### 3.3. Pre-processing and Connectome construction

FMRI data were pre-processed according to a standard pipeline: motion correction, mean intensity subtraction, passband filtering with cutoff frequencies of [0.005-0.1 Hz] and skull removal. To account for potential noise from physiological processes such as cardiac and respiratory fluctuations, eight covariates of no interest were identified for inclusion in our analyses as nuisance variables (Saad et al., 2013). For this aim, regressors for white matter (WM), cerebrospinal fluid (CSF), and the 6 motion parameters for each individual were estimated. Spatial smoothing was performed only by averaging the voxels for the ROIs. Global signal was not regressed to avoid spurious anticorrelations (Murphy and Fox, 2017). To further reduce the effects of motion, correction for frame-wise displacement was carried out as described in Power et al. (2012). For both datasets, the DWI data were cleaned by eddy current interference, and the skull has been removed. Linear registration has been applied between the Power atlas (Power et al., 2011) and the T1 reference volume by using linear registration with 12 degrees of freedom. The Power’s atlas defines 264 non-overlapping functional ROIs. Each ROI is a 5mm sphere around a peak voxel of significant activation during cognitive tasks calculate with a meta-analytic approach on the neuroimaging literature on this topic. This atlas is to date considered one of the most accurate for functional mapping (as shown e.g. in Gordon et al. (2014)).

Tractographies for all subjects were generated processing DWI data with the Python library Dipy (Garyfallidis et al., 2014). In particular, we used the constant solid angle Q-ball model, and a deterministic algorithm called Euler Delta Crossings was used stemming from 2,000,000 seed-points and stopping when the fractional anisotropy was smaller than *<* 0.05. This low threshold was chosen to allow the fibers to enter the small ROIs of the chosen atlas. Tracts shorter than 30 mm or in which a sharp angle (larger than 75°) occurred were discarded. It is important to mention that the choice of more common parameters, as an angle threshold of 60° or fractional anisotropy threshold of 0.2, would not allow in the used data the fiber bundles from the posterior cingulate to enter the prefrontal cortex, compromising the DMN analysis. The final result yielded about 250,000 fibers. To construct the connectome, the graph nodes were determined using the 264 regions in the Power atlas. Specifically, the structural connectome was built counting the number of tracts connecting two regions, for any pair of regions. The same regions were used to compute the functional activity just averaging the voxel activity in each area.

### 3.4. Structurally constrained Autoregressive model

A multivariate autoregressive model of order *n* (MAR(*n*)) is a stochastic process defining *r* variables **y**(*t*) as linearly dependent on their own previous values and on a stochastic term:

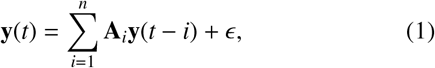

where the *r*-by-*r* coefficient matrices **A**_*i*_ are the model parameters and *E* is the additive Gaussian noise. The weight matrices **A**_*i*_ describes the linear dependencies between the *r* timeseries [*y*_1_*,…, y_r_*]^*T*^; therefore, they can be intended as a functional connectivity matrix at different time lags. The subscript *i* indicates the order or the time lag. The aim of the proposed method is to infer a functional connectivity justified b y t he structural connectivity, so that the functional activity of the brain can be described only using the direct physical connections between the brain regions. To obtain this result, we resort to a set of constraints on the above MAR model, which force the MAR model to only fit the parameters associated to existing structural connections. In this way the signal in a brain region is described by a linear combination of the function of structurally connected regions only:

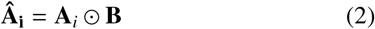

where ⊙ denotes the Hadamard or element-wise product, and **B** is an indicator matrix of the structural connectivity **A**_init_, defined as

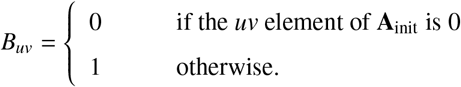

This **B** matrix is used as a structural bias into the model, constraining each **A**_*i*_ to have zero elements where no structural connectivity appears. Given an initial structural connectivity matrix **A**_init_ and the functional signal in a column-vector **y** for all *T* time-frames, an effective connectivity matrix can be determined minimizing a reconstruction error of the MAR model in eq. (1) subject to the structural constraint in eq. (2) as follows:

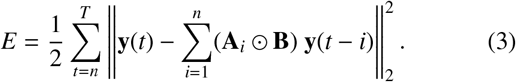

The effective connectivity matrix is therefore the matrix **Â**_*i*_ that minimizes the reconstruction error above. The choice of the model order *n* can be defined by Akaike information criterion (Akaike, 1974), or empirically in cross-validation settings. In our experiments we adopted this latter approach to choose the order of the CMAR model. As observed empirically, relaxing the constraints, which we introduced in our method, was leading to lower prediction error, though it was introducing structural connections which were not biologically plausible.

However, the model described so far can only relate direct structural connections to functional series. To overcome this limitation, equation (3) is modified including propagators that iterate through several steps through the connectome inspired by the neural field theory propagators (Robinson, 2012). According to the propagator formulation of neural interaction introduced by Robinson (2012), firing neural rate within a brain network **Q** can be computed as the sum of external electrical activity **N** and by the firing rates coming from other neurons. Those rates can be directly evoked or incorporate activity coming from synaptic path **Λ**: **Q** = Λ**Q** + **N**. More specifically, the *m* steps of propagation can be defined by the sum **Λ = Λ**^(**0**)^ + **Λ**^(**2**)^ + **Λ**^(**3**)^ + … + **Λ**^(**m**)^ (following the notation of Robinson (2012)). An example of this propagation from one region to another of the brain is depicted in Figure 1. In our implementation we neglected the external input **N**. Conversely to partial correlation (Zalesky et al., 2012), this multi-step formulation is based on physical connections. Despite the temporal resolution difference between firing rate and BOLD signal, same assumption of propagation can be made for the BOLD signal case given the relationship to neuronal stimulus (Drysdale et al., 2010). This will also be comparable to the multisteps in causal graphs (Chicharro and Panzeri, 2014), to connectome spreading dynamics (Mišić et al., 2015), and to the formulation of indirect Granger causality (Dhamala et al., 2008). Given those tools, equation (3) can be reformulated taking into account higher-order effects as downstream effects of the direct interactions. For instance, for the first order of indirect connections the reconstruction error becomes

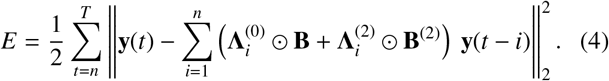

In practice, 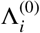 comprises the coefficients of effective connectivity related to direct connections, and it is initially set as the structural connectivity matrix **A**_*i*_. A further coefficient matrix is introduced to weight the indirect connection of first order, and the matrix **B**^(2)^ comprises the indirect structural connections of first order likewise **B** for the direct connections. For low resolution networks **B**^(2)^ can be easily given by the complement of **B**, since the first order indirect connections will cover the remaining brain connections. The reason why the structural matrix is used as initial guess, it is only given by the fact that it is believed to be a decent initial guess in the gradient descent context, though there are no arguments against using other values including random estimates. To obtain the effective connectivity matrix we first fit the model parameters by using a gradient descent approach. The direction of the gradient can be computed for all matrices **A**_*j*_ of a specified order for the direct connections as follows:

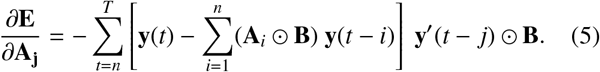

and the update rule is therefore **A**_*j*_ = (**A**_*j*_ + *η* · *∂***E***/∂***A**_*j*_), where *η* is the learning rate. For the multistep case **Λ**_*i*_ the gradient direction can be computed deriving according to the individual **Λ**_**i**_, leading an expression similar to (5). In the reported experiments we used 5000 iterations during the gradient descent, and a learning rate *η* = 3 · 10^−5^. According to our model, the initial condition is limited by the structural connectivity matrix **A**_init_, likewise for the indirect case. We tried different normalizations of the matrix (as binarization and division for the maximum value) and the same minimum was reached, even if different speed of convergence has been observed. A further improvement is given by deconvolving the BOLD signal to use a more appropriate neural response. Assuming a common HRF is shared across the various spontaneous point process events at a given voxel, the BOLD signal can be seen as the result of the convolution of neural states *s*(*t*) and HRF *h*(*t*):

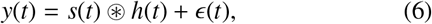

where *ϵ*(*t*) represents additive noise. Once calculated *h*(*t*), we can obtain an approximation of the neural signal *s*(*t*) from the observed data using a Wiener filter (Wu et al., 2013). We conducted our experiments first by using the initial preprocessed BOLD time series and then again using the estimated neural response after the deconvolution 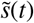. In this view, the previous optimizations should be seen substituting *y*(*t*) with *s*(*t*).

Once optimal effective connectivity matrices **A**_*i*_ for each subject are obtained, Granger causality can be computed defining the connectome as a directed graph. Several ways to compute Granger causality both in the frequency and time domain were proposed. The most known is the reduced model (Geweke, 1984), where error of linear dependence for two multiple time series is measured quantifying the causality of one against the other. As shown by Stokes and Purdon (2017), fitting a reduced model for spectral Granger causality can lead to a strong bias or very large variability depending on the choice of the order. Consequently, we computed the Granger causality matrix representing causality given the ratio of residuals (prediction error using or not a driver ROI), by using the code related to the manuscript of Luo et al. (2013).

### 3.5. Simulations and Default Mode Network Analysis

For the simulations we used the same ground-truth and settings of the reference paper for the NetSim, a well-known evaluation of network modelling methods for fMRI (Smith et al., 2011). Briefly, we used 50 simulations given by 5 nodes/processes (*Y*1*, Y*2*, Y*3*, Y*4*, Y*5) 10 minutes long and obtained as described in Smith et al. (2011), defined by using repetition time 3s and standard deviation of the haemodynamic response function (HRF) 0.5s. Figure 3 (a) depicts the defined ground-truth causal inferences.

The DMN analysis was used to prove that the proposed method produces sensible results in line with current literature. More specifically, in this experiments starting from a whole brain structural connectivity and related resting-state fMRI series, an effective connectivity was constructed, and then the specific effective connections were analyzed. Moreover, the proposed method was compared to DCM considering all possible causality directions. The optimal DCM was obtained according to the Bayesian information criterion (BIC). Tractography was chosen as a prior for sigmoidal model of the DCM, as suggested by Stephan et al. (2009) being a good estimate of the prior, though other configurations are also possible. We considered indistinctly excitatory and inhibitory causality jointly. Further limitations of the DCM were related to the fact that resting-state does not have a specific driving input as in other DCM models (Adams et al., 2013). We overcame this limitation by using a stochastic DCM (Daunizeau et al., 2014) that adds time-dependent fluctuations in neuronal states.

### 3.6. Classification of Autism Spectrum Disorder and Typically Developing subjects

To test the descriptive power of the effective connectivity computed by using the proposed approach, we carried out a classification task discriminating between ASD and TD individuals, using all three types of connectivity: functional, structural and effective. We investigated the most significant connections obtained through the weights of a trained support vector machine (SVM) (Crimi et al., 2017). Those were the weights larger than the 95^*th*^ percentile or smaller than the 5^*th*^ percentile of a random weight distribution which represents the null hypothesis. The null hypothesis SVM weights were obtained by performing 1000 random permutations of the labels of the two groups.

### 3.7. Effective Brain Community Detection

To assess whether the proposed method produces effective connectivity information characterizing the structural connectivity enriched with functional information, a community detection analysis was performed using a group-wise graph clustering algorithm recently proposed in Crimi et al. (2016) both on the set of structural connectivity matrices and on the effective connectivity matrices.

Given a set of connectivity matrices 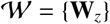 representing undirected weighted graphs with positive weights, each normalized graph Laplacian is built as 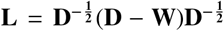, where **D** is the diagonal degree matrix of **W**. However, in general, the connectivity matrices resulting from the above CMAR model computed for each subject are asymmetric (i.e., edges are directed), thus, they were converted to undirected graphs aiming at maintaining the properties of the original graphs estimated from CMAR. To this aim, a symmetrization based on random walk was applied (Malliaros and Vazirgiannis, 2013).

More specifically, given a directed graph **M**, the transition matrix of the random walk can be defined as 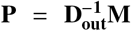, where **D_out_** is a diagonal matrix built using nodes’ out-degree. The symmetric graph can be therefore defined as 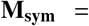 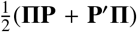, where Π is a the diagonal matrix that defines the probability of a walker to stay in each node in a stationary distribution, defined as 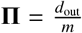, where *d_out_* is the vector of the out-degree of each node and *m* is the number of nodes. Thank to this new representation, the pipeline described by the authors in Crimi et al. (2016) can be applied, generating the normalized graph Laplacians for each subject, performing the joint diagonalization of multiple Laplacians to find a unique eigenspace and, finally, applying spectral clustering on the smallest joint eigenvectors.

In order to decide the number of clusters, as usual in spectral clustering, we looked at the spectral gap of the mean approximated eigenvalues. Supplementary Figure 2 in the supporting information depicts mean approximated eigenvalues where a gap between the 8th and 9th values is visible. This was assessed both visually and computationally. The cluster functional separation (CFS) among the brain nodes was defined as the average ratio between the intr- and inter-cluster cross-correlation. as

follows:

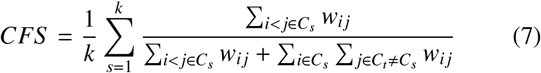

where *w_ij_* is the functional cross-correlation of the time-series for nodes *i* and *j*.

## 4. Results

The validation of the proposed method has been performed first by using simulations and by looking in a subset of areas related to the DMN, then a classification task (Crimi et al., 2017) aiming at discriminating autistic and typically developing children has been carried out, and lastly brain community detection framework has been used jointly to the proposed model. When possible, a comparison to the well-known DCM approach was carried out. The DMN and effective community experiments were performed by using the diffusion tensor and fMRI volumes of 200 healthy subjects from the Nathan Kline Institute-Rockland dataset (Nooner et al., 2012). The classification experiment has been carried out from data of the ABIDE-II dataset (Fishman et al., 2015).

### 4.1. Structurally Constrained Autoregressive Model

Figure 2 depicts an example of initial structural tractogram and final effective connectivity diagram, given by chord diagrams where each node represent a ROI. In this figure it is shown that some connections are canceled out resulting in a less dense chord diagram,Those results are for the case considering both direct and indirect connections, respectively 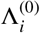 and 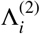 in Figure 1. Although not clearly visible when comparing visually adjacency matrices, the initial structural connectome had mean binary connections *μ_s_* = 8198 against the mean binary effective connections *μ_e_* = 3169 across the whole dataset. Comparing the proposed approach when using only the direct structural connections and when considering also the indirect connections, it was noted that using the latter was producing a lower reconstruction error though most of the influence is already given by the direct connections, as shown in Figure 5 (a). In our experiments we define reconstruction error as the difference between a predicted and real functional series given a connectivity matrix and the value of a previous time point (as in equation (3) and (4)). We optimized the weights for the direct and indirect connections separately. Namely, we obtain the coefficients first for the direct connections and then for the indirect connections. This was done since a parallel optimization was shown to be slower due to the fact that the two processes compete with each other. Lastly, by using the estimated neural signals instead of the BOLD signal further improvements are obtainefd as shown in Figure 6.

**Figure 2:**
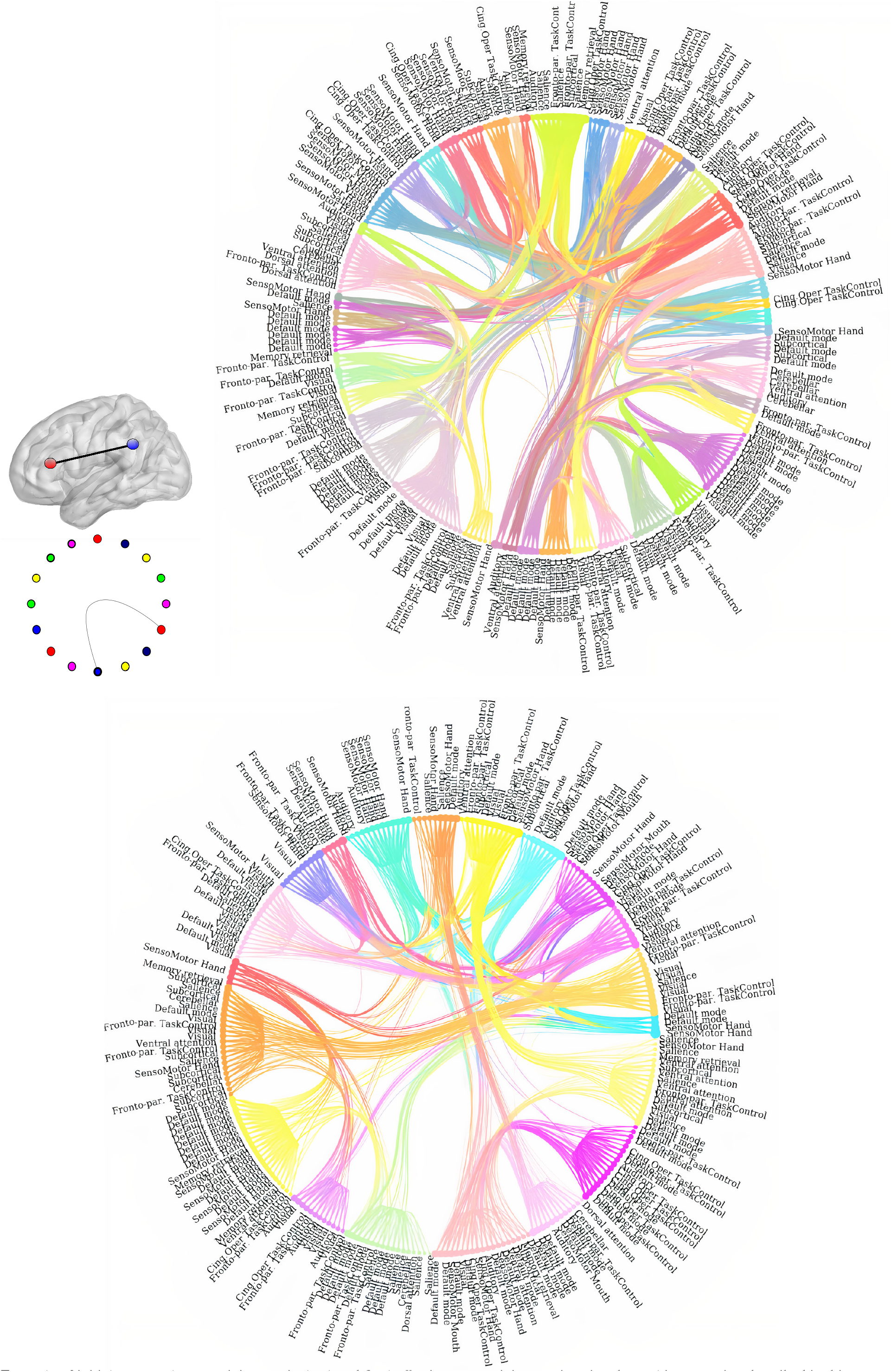
Example of initial structural connectivity matrix (top) and final effective connectivity matrix using the multistep version described in this paper (bottom) for the same subject, as chord diagrams for the 264 ROIs (Power et al., 2011) in which the brain is partitioned, with visibly a less dense diagram for the final effective connectivity. Color-code does not represent ROIs but connections between them. The ROIs defined as uncertain in (Power et al., 2011) are not labeled.

Figure 5 (b) highlights the beneficial effect of using more time points in the MAR model for the autism dataset (Fishman et al., 2015). As expected, the higher the order of MAR model the better was the reconstruction error of the signal, though at some point the improvements were not worthwhile. In particular, it was noted that the reconstruction error of MAR(2) is always significantly smaller than MAR(1). However, in all our experiments, going beyond order 2 did not improve the reconstruction error compared to MAR(2). This is probably related to the type of signal. Indeed BOLD signal in used fMRI recordings has a sampling rate that compared to the underlying brain activity makes it useless to go beyond two time steps.

Once the final effective connectivity matrix is computed, it can be used to quantify the Granger causality. Supplementary Figure 1 in the Supporting Information depicts the average of the resulting autocovariance matrices of the Granger causality, where non-zero elements represent causality from one region to another. As for the resulting effective connectivity matrix, in this matrix some regions have a strong influence compared to other, and some regions of the same network influence each other. Some expected asymmetry in the resulting matrices was also observed. The identified causalities appeared as a subset of the effective connectivity detected by the proposed model. As empirically observed, removing the structural constraint and in lieu using sparsity regularization (Valdés-Sosa et al., 2005) was leading to a lower reconstruction error, but it was introducing connections that were anatomically implausible.

### 4.2. Simulations and Default Mode Network analysis

After initial analysis on reconstruction error, the first experiments were performed by using simulations and real data but limited to the DMN. The results were compared to those obtained by using a physiologically defined model in contrast to the proposed data-driven model: the resting-state DCM (Daunizeau et al., 2014)

For the simulations we used the same ground-truth and settings of the reference paper for the NetSim, a well-known evaluation of network modelling methods for fMRI (Smith et al., 2011). Briefly, reproducing Smith et al. (Smith et al., 2011), we used the simulations given by 5 temporal processes (*Y*1*, Y*2*, Y*3*, Y*4*, Y*5) 10 minutes long. Those processes are related by causal inferences represented by the directed graph shown in Figure 3 (a). The generated time series and the connectivity matrix without causal information are then used by the proposed algorithm (only direct connections and first order), and by the optimal DCM according to the Bayesian information criterion. The difference between the estimated causal inferences and the ground-truth are shown in Figure 3 (b). Difference was given by the number of causal directions that were quantified wrongly. If one algorithm was estimating only opposite direction than the ground-truth, the difference was quantified as ‘2’. If for one edge one method estimated bidirectional causality but only one was correct, the difference was quantified as ‘1’. It was noted that for both the proposed model and DCM the causal inference *Y*1 → *Y*2 was generally estimated incorrectly but the DCM was having even more pronounced differences with the ground-truth. Comparing the two models by using t-test the mismatch had a statistical difference of t-score of 1.75.

**Figure 3:**
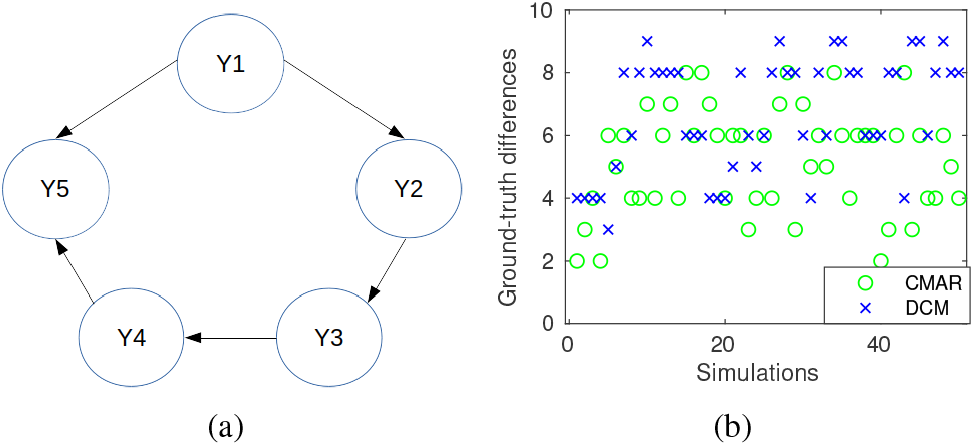
(a) Ground-truth simulated causalities. (b) Difference between the ground-truth and the estimated causal matrix by the proposed method and DCM for 50 independent simulations, each simulation mismatch reported along the x-axis. For each simulation we reported the mismatch per model, and it was visible that DCM had more mismatch from the preferred model, the higher the data point, the more mismatch the model produced.

Afterward, we assessed whether the most important structural connections relating the functional areas of DMN were maintained after the optimization of the connections with our CMAR model and Granger causality. For this purpose, we analyzed a subset of DMN areas and the connections that relate each other. We focused on some DMN regions that are not directly structurally connected, though they show some activity simultaneously (Greicius et al., 2003): frontal middle gyrus, posterior cingulate cortex, and fusiform gyrus; all both left and right. The proposed model has been used on a whole dataset by using these areas showing that they persist after the autoregressive model, and still identify areas that are physically connected and used functionally as shown in the example in Figure 4(a). The resulting effective connectivity DMN was compared for completeness to the optimal DCM that was selected among all the models having all possible direction combinations. By neglecting the commisural connections, the possible models to be tested were 27. We report the selected optimal DCM is depicted in Figure 4 (b). An example of tractography among those ROIs for one subject is shown in Figure 4 (c). Carrying out a brain-wide CMAR optimization, the relevant DMN connections were maintained. Each DMN region before the CMAR had on average about 30 connections (not depicted in the graph to avoid cluttering), while after the CMAR the average connections were 8 and those were the most relevant for the DMN regions based upon literature Greicius et al. (2003). Conversely, the DCM needed to have the 6 used regions specified before hand. Moreover, the CMAR/Granger analysis did not erase the relavant connections, and found bidirectional causality between the cingulate and prefrontal cortex, though the causalities from the posterior cingolum to the prefrontal cortex were stronger. While for the selected DCM the stronger causalities were in the opposite direction. It is worthwhile to mention that all DCMs required specific settings discussed in the Discussion and Methods sections.

**Figure 4:**
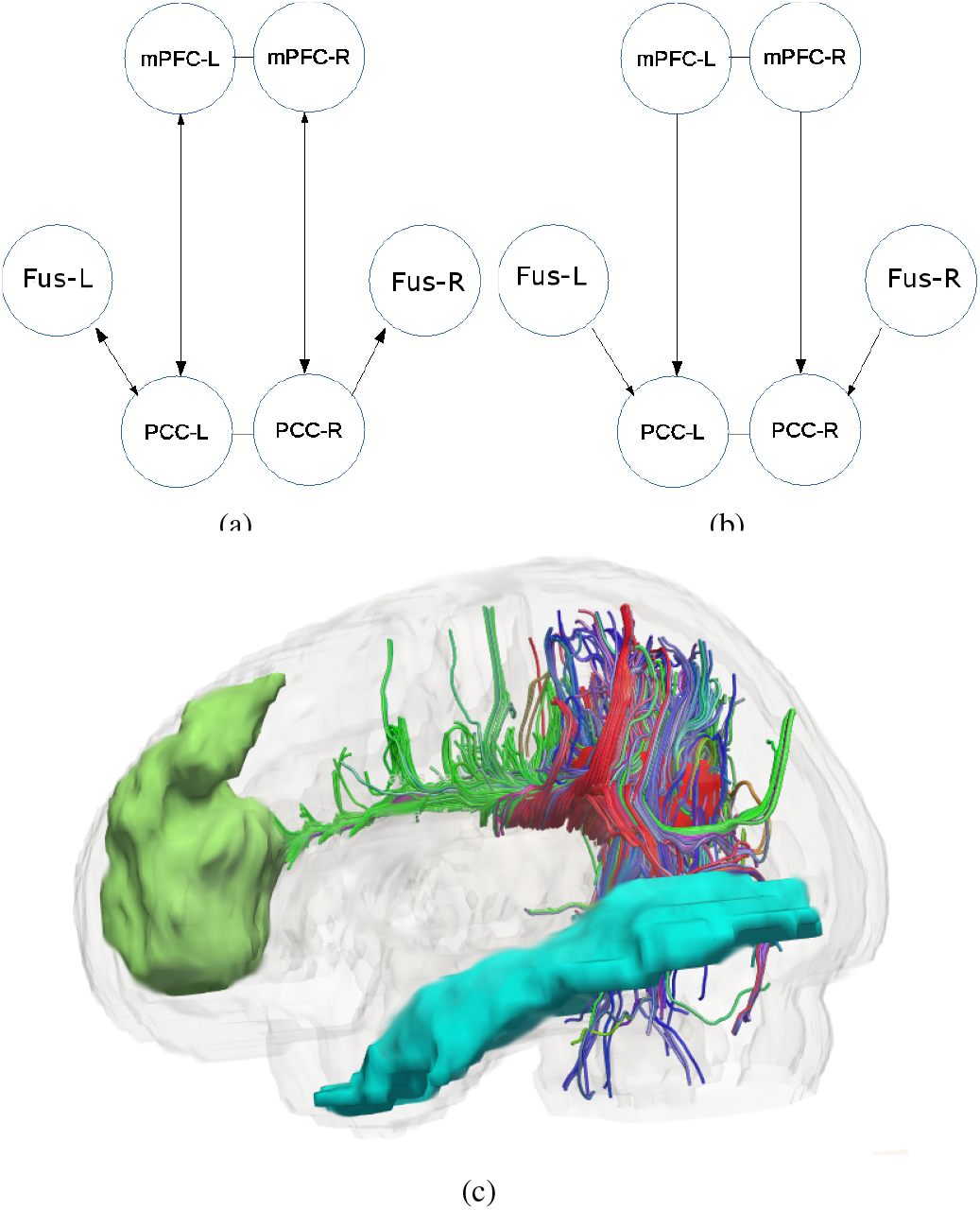
(a) and (b) are respective graphical representations of the resulting Granger causality and DCM for the DMN experiment. The reported acronyms are ‘mPFC’ = medial PreFrontal Cortex, ‘PCC’ = Posterior Cingulate Cortex, ‘L’=Left, and ‘R’=Right. This reflects structural and functional expected connections depicted in (c): Sagittal example of structural tracts connecting the functional regions of interest selected as related to the DMN. Those ROIs are obtained as a subset of the DMN ROIs of the Power atlas according to the coordinates specified in (Shirer et al., 2012), if necessary more than one ROI of the Power atlas was merged. The PCCs are depicted as red, mPFCs are depicted as green and the fusiform gyri in cyan.

### 4.3. Classification with Effective Connectivity in Autism

A classification task between ASD and TD individuals was used to further prove the descriptive power of the effective connectivity computed using the proposed approach.

We investigated the most significant connections obtained through the weights of a trained support vector machine (SVM). Those were the weights larger than the 95^*th*^ percentile or smaller than the 5^*th*^ percentile of a random weight distribution which represents the null hypothesis, as depicted in Figure 7 (d-f). Of note, the most discriminating connections of the MAR model were very different from those determined with struc tural connectomes. Figure 7 (d-f) shows the whole spectrum for the averaged SVM weights respectively for the structural, effective MAR(1) and functional connectomes. It is noticeable that the weights for the MAR cases are larger than those related to the original structural connections, and the weights related to the functional data are more numerous. Given the fact the functional connectivity can be given by anticorrelations, positive and negative weights are present. The detected connections using the structural data comprised ipsilaterally spread connections in agreement with previous studies (Li et al., 2014; Zhang et al., 2017). The detected functional connections also confirmed previous results obtained by using another dataset (Yahata et al., 2016). For instance, the connections between the left frontal pole and the right central gyrus, as well as the left inferior frontal gyrus and the right inferior temporal gyrus, were also identified as significantly different between cases and controls by our model. The detected effective connections partially resemble the detected structural and the functional connections. Interestingly, these detected features are even more in agreement with the discriminant features identified through previous studies: as such the ectopic connectivity of precentral and striatal and occipital regions (Noonan et al., 2009), the left superior parietal and right inferior frontal gyrus, and rectus and cuneus region (Yahata et al., 2016). The classification accuracy in discriminating ASD from TD subjects by using the identified connections was respectively for the structural, functional, and effective connectivity using MAR(1) and MAR(2) 60.57%, 69.1%, 70.21%, and 72.34%. This showed the efficacy of using effective connectivity matrices instead of others for the proposed experiment, although the experiment should be repeated with larger datasets. Lastly, it was noted that the detected functional connections were larger in number and both negative and positive to compensate the signs given by correlations and anticorrelation.

**Figure 5:**
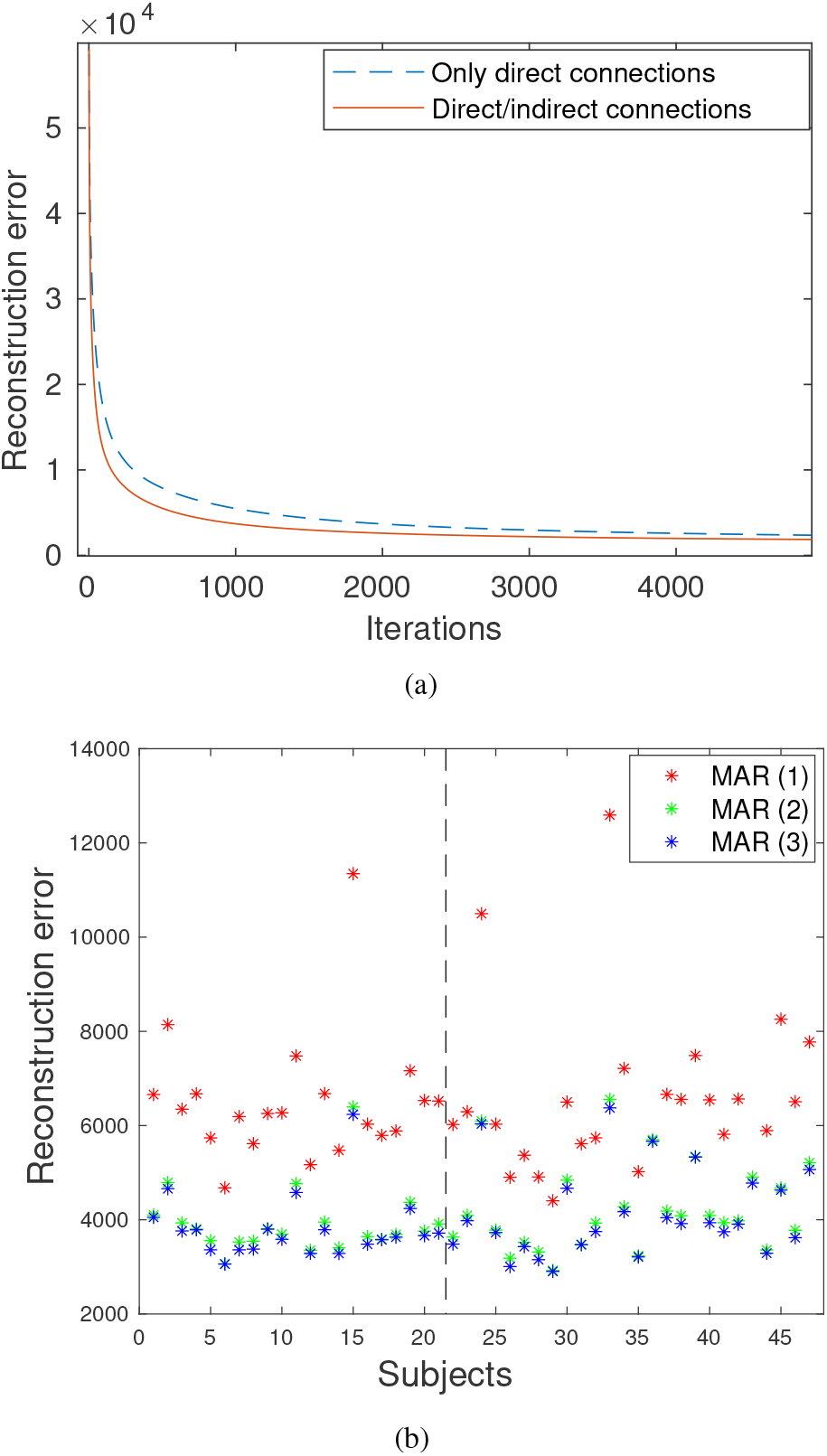
(a) Example of reconstruction error for one subject of the Nathan Kline Institute-Rockland dataset by using only direct structural connections and direct jointly to indirect connections. (b) Final reconstruction errors for different orders using the ABIDE-II dataset. The stars at the left and right of the line depict respectively the errors for the control (TD) and case (ASD) subjects from the classification experiment.

**Figure 6:**
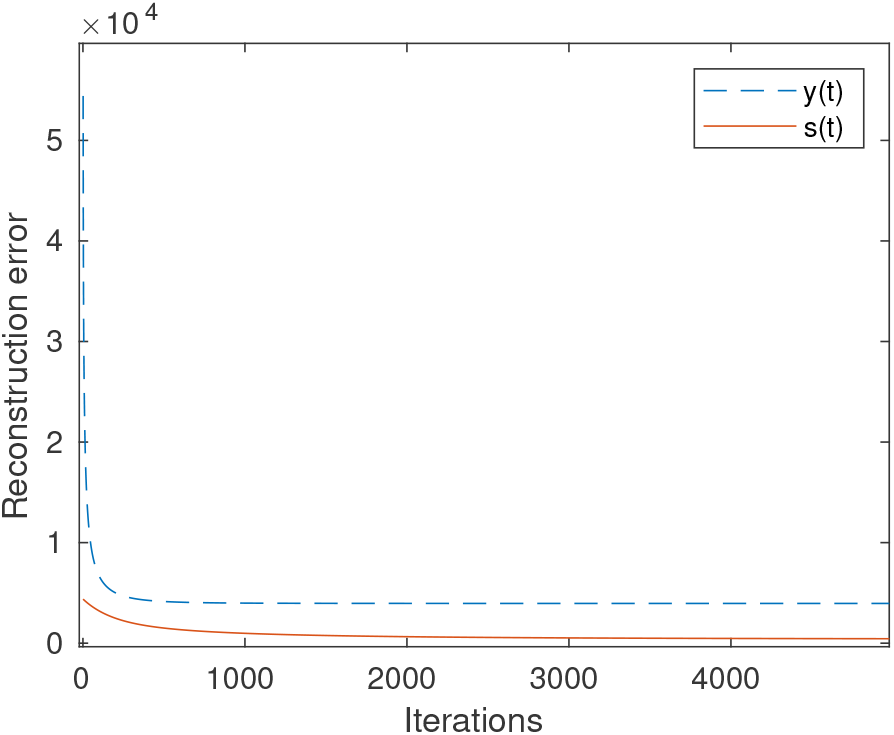
Example of reconstruction error by using the preprocessed BOLD *y*(*t*) and the estimated neural signal s(y). Here, the deconvolution is performed before carrying out the MAR analysis.

**Figure 7:**
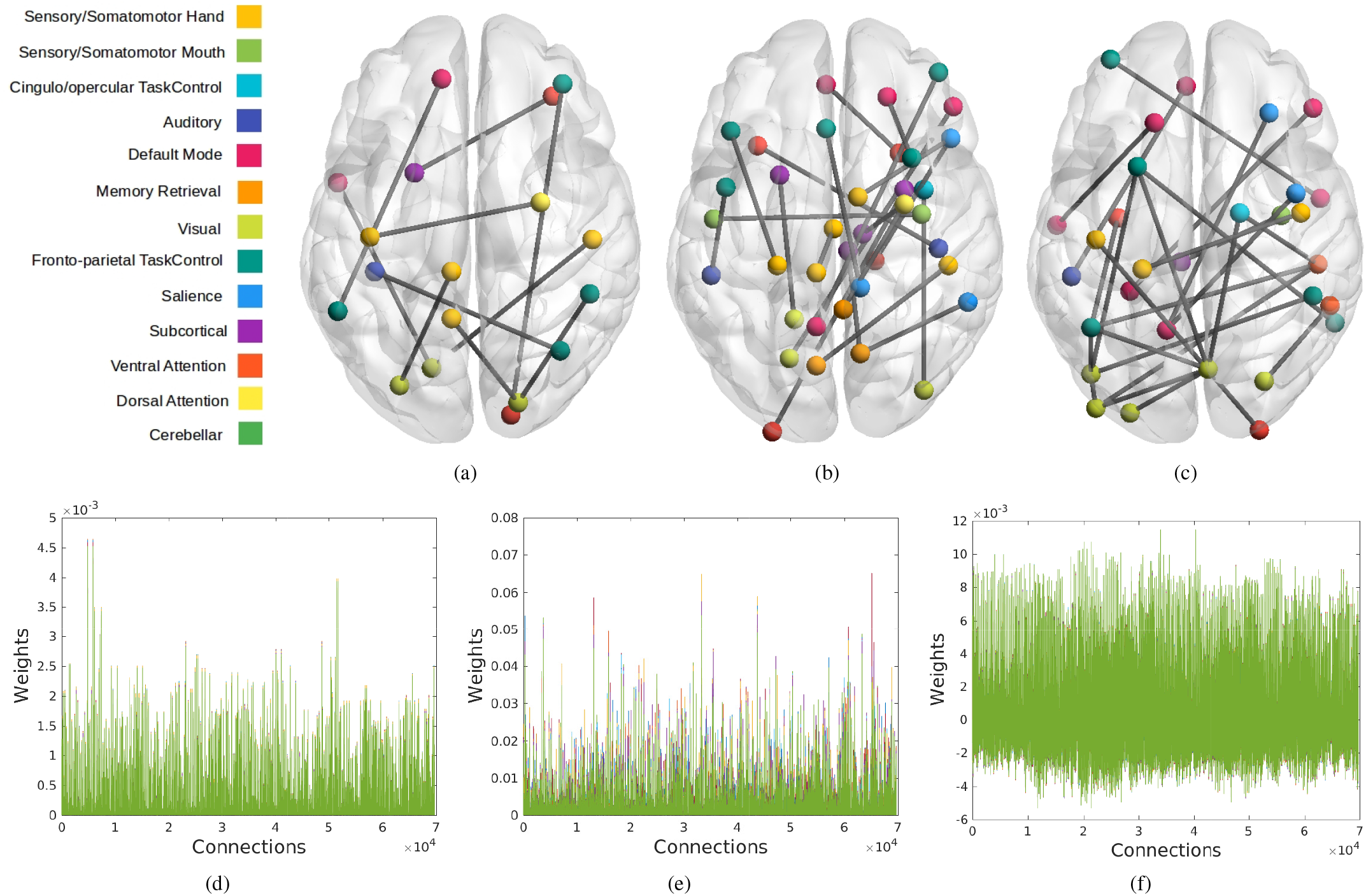
Axial views of most discriminant connections according to the SVM weights (a) original connectomes, (b) MAR(1) effective connectomes, and (c) functional connectomes. (d), (e) and (f) show the average weights respectively for the structural, the MAR(1) effective and functional connectomes. The highlighted structural connections in (a) are mostly ipsilateral and few commisural connections. Most of the detected functional connections (c) are already known (Yahata et al., 2016) as the link between the left frontal pole and the right central gyrus, and the left inferior frontal gyrus and the right inferior temporal gyrus. The effective connections in (b) also highlight known connections as the left superior parietal and right Inferior frontal gyrus, and rectus and cuneus region (Yahata et al., 2016).

### 4.4. Effective Brain Community Detection

This latter experiment aimed at identifying community in the effective connectome which better represent the original functional signal in terms of segregation. By analyzing the groupwise eigenvalues resulting from the joint Laplacians diagonalization, a spectral gap was noted at the 4^*th*^ and 8^*th*^ eigenvalues for both structural and effective connectivity matrices, in agreement with previous studies on other datasets (Hagmann et al., 2008) and as depicted in supplementary Figure 2 (Supporting Inforamtion). The value *k* = 4 has been neglected since the clustering for *k* = 4 would have led to a simplistic representation of the brain. Therefore *k* = 8 has been used. The resulting clustering of the brain regions based on the structural connectome (Figure 8 (a)) and on the effective connectome (Figure 8 (b)) are fewer while preserving the overall organization. The reconstruction error shown per subject in Figure 8 (c) indicates, as expected, that the lowest error is given when considering the whole network in the CMAR computation.

**Figure 8:**
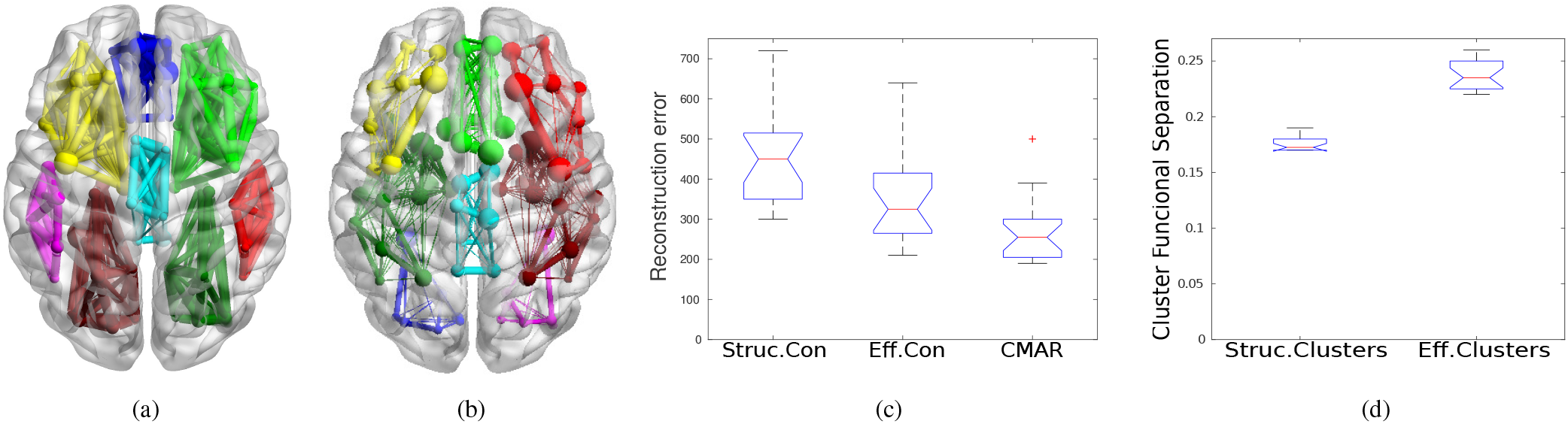
Axial view of joint spectral clustering using k=8 is depicted in (a) for the original structural joint eigenspace, and (b) on the joint eigenspace given by effective connectivity matrices. The resulting connectivity matrices as shown in Figure 2 (b) and the related clustering in this subfigure (d) are not only sparser than the original, they also lead to better reconstruction of the functional data as shown in the other figures: Subfigure (c) depicts the reconstruction error of CMAR model (represented as errorbars) after converging to the effective connectivity (green circles) or according to block-wise MAR based on the structural communities and the effective communities. The lower the better. Subfigure (d) depicts the functional segregation of clusters using the effective communities or structural communities represented as errorbars. The higher the better.

Evidence was also obtained when analyzing the cluster functional separation, defined as the average ratio between the intra- and inter-cluster cross-correlation (Crimi et al., 2016). This index has been computed for both structural and effective clustering results. Figure 8 (d) indicates that cluster functional separation of clusters determined using our CMAR approach is significantly higher when compared with the structural clusters (*p <* 0.001) according to converted t-tests, demonstrating that the effective clusters are also underpinned by the functional connectivity. More specifically, we also devised an analysis to assess whether the clusters obtained from the autoregressive modeled data are more meaningful in relation to the fMRI timeseries than the clusters obtained from the structural information. To this end, we carried out a block-wise definition of the effective connectivity matrices where one block at a time, defined by the brain regions belonging to a cluster, was used in a CMAR model involving only the relative fMRI series. Then the reconstruction error of the fitted CMAR models for each cluster was summed up over all clusters and compared to each other. The underlying intuition is that partitioning the brain using an effective connectivity information would remove those structural connections that are meaningless from a functional perspective, at least in the analyzed experimental data. This is the overall message conveyed in Figure 8.

## 5. Discussion

Investigation of how structure influence function is present in several fields. For example, how the function of a protein can be predicted by its sequence and structure is a common issue in proteomics (Lee et al., 2007; Watson et al., 2005). In neuroscience, it is now well established that functional segregation is a basic principle of brain organization (Hermundstad et al., 2013). However, this principle is difficult to prove and the easiest approach is to investigate it through functional connectivity analysis, which is usually based on the determination of functional correlations between brain regions. Causal influence is most likely the key to deliver insights in neuroscience and physiology (Bielczyk et al., 2018). In this context, the proposed model combines structural and functional information with the aim of defining effective connectivity. Initially, given the initial brain-wide structural connection the autoregressive model acts as a filter removing the connections that are not used in a related functional series, then causality is investigated according to Granger causality. The previous experiments showed that a resulting connectivity matrix can be used to relatively predict BOLD signal. The use of indirect connections, rather than only direct connections, showed a lower reconstruction error in predicting the BOLD signal though a slower convergence, though this was a slight improvement. As described in previous studies mapping brain functional connectivity from diffusion imaging suggested Becker et al. (2018), the propagation of neural signals through direct and short structural walks has the strongest influence in the resulting functional connectivity, and this can be true also for our case. The comparison between using BOLD signal and estimated neural signal further shows the importance of isolating the HRF from BOLD signal.

In the simulation experiments the proposed CMAR estimated causal inferences with lower difference from the ground-truth compared to the DCM. The differences of the CMAR from the ground-truth were mainly in associating opposite causal inference between the node *Y*1 and *Y*2 compared to the ground-truth. The DCM had even more mismatched causal inferences. In the experiments with the DMN regions, the proposed model starting from brain-wide connection, it was able to identify the relevant connections of the DMN without removing them. The causalities determined through the CMAR were partially overlapping with those identified by DCM, without the extensive design of the models described below. Without a proper groundtruth it is difficult to establish whether the data-driven or the selected DCM model produced a more correct estimation of the causalities. In fact, in previous studies DCM applied to DMN has produced inconsistent results (Razi et al., 2015). A further hurdle using the DCM was given by the need of specifying a driving input, which is not clearly defined for the resting-state case (Adams et al., 2013). Nevertheless, the CMAR was able to extract from all the given connections meaningful DMN networks while the DCM required the investigation of all possible causal directions, with the limitation of increasing complexity increasing the number of nodes. This shows that Granger causality combined with the proposed CMAR model is a valid explorative data-driven approach. Yet DCM is mostly a validation tool, and they should be seen as complementary. Still, our aim is not to solve all possible controversies related to causality, especially considering the criticisms that those models are more temporal correlations (Bielczyk et al., 2018; Etkin, 2018), but to provide a straightforward method to identify structural connections effectively used during tasks or rest that can be used for biomarkers or other purposes.

The experiment of classification between autistic and typically developing subjects also improved if the resulting effective connections were used. This led to the hypothesis that using features from the resulting effective connectivity removes those structural connections that were also meaningless from a functional perspective or combine both features, leading to a better performance in terms of clustering or classification. Indeed the classification was better if effective connectivity was used instead of the structural or functional connectivity. Moreover, the identified discriminant connections were in line with connections detected by other works described in the Results section (Yahata et al., 2016; Mastrovito et al., 2018). Although the improvement obtained in this experiment should be repeated with larger sample size, the hypothesis is that using features obtained with our model offers the advantage of combining functional and structural features, with advantages compared on using only one type of features.

In our experiments, the identification of the effectively used structural connections had clearly an impact on the subsequent clustering, which is reflected in Figure 8 (c-d). Namely, the CMAR while preserving the structural partitioning, also optimized the graph cut minimizing the loss of functional interactions. In fact, when removing some connections according to the clustering results, the communities determined from the effective connectivity matrix appeared to be more self explanatory in terms of functional activity than the communities obtained from structural connectivity alone.

In all our experiments, contrary to DCM that requires a detailed setup of numerous experimental setup, the complexity of the proposed method was limited to the first order of the autoregressive model and of the propagators. The former takes into account the number of prior time points within the BOLD series that need to be examined, the latter the number of nodes within the structural connectome that have to be crossed (steps). The autoregressive order was found empirically to be optimal at 2, while the steps of the propagator were also limited at 2. The low values for the autoregressive model are most likely related to low temporal resolution of the BOLD signal. Regarding instead the propagation through indirect connections, due to the resolution of the atlas the used matrix **B**^(**2**)^ was already comprising all complement of the matrix **B**. Once again, most likely due to the nature of the BOLD signal, the strongest influence was given by the direct connections as shown in the example in Figure 5.

Besides those observations, CMAR is independent on whether the BOLD series is task-based or resting-state. Deconvolving the signal before the use in the MAR model is a useful extension to the proposed model which can produce more accurate estimates effective connectivity. The proposed approach can be seen as similar to previous works relating precision matrix from functional series to structural information (Deligianni et al., 2013; Hinne et al., 2014; Ng et al., 2012), but where those models constrain functional connectivity, the proposed model used it to define effective connectivity.

Some studies have pointed out that functional connections can be violated from the underlying structural connections (Preti and Van De Ville, 2019; Medaglia et al., 2018; Tyszka et al., 2011) and this can have an impact in the prediction. Those violations can arise either by the differences between effective connectivity and functional connectivity (Goldenberg and Galvaán, 2015), or limitations of the diffusion imaging acquisitions. Effective connectivity captures the biophysical transfer of activity between brain areas along structural pathways, it fundamentally differs from functional connectivity which only manifests the measurable consequences of such interactions, described as correlations. Moreover, the nature of the DWI signal can estimate fiber density, but might be inaccurate to capture the true synaptic interaction strength between areas, especially considering short-term plasticity, which may alter the interaction strength especially during a task setting (Hahn et al., 2019). Despite those discrepancies, relationships between functional and structural connectivity exists. A further example of this is the impact of corpus callosus agenesis, which for example can reduce the underlying structural connections related to the DMN to the cingolum, affecting the functional connectivity (Hearne et al., 2019). Given these observations the proposed method is more focused on the effective connectivity capturing biophysical transfer of activity between brain areas, and this limitation cannot unravel all possible aspects of functional connectivity, which are yet to be understood.

Summarizing, despite the limited temporal resolution of BOLD signals, models such as those defined by the Granger causality can make explicit the definition of causality by the use of MAR models that are easy to validate (Goebel et al., 2003). The effective connectivity inferred by the proposed constrained MAR model highlights a different brain architecture underpinned by both structural and functional connectivity, which is in-line with current neuroscience principles (Passingham et al., 2002). This represents a valid alternative to other approaches which are computationally limited.

The DMN experiment served as a proof of concept of the validity of the method, conveying results in line with known connections from literature (Demertzi et al.; Greicius et al., 2003). Other experiments highlighted also the benefit of this approach. Namely, the effective influence can be identified among specific brain regions with which they interact, given by effective connectivity rather than just functional co-activation or structural connection (Sridharan et al., 2008). This method can lead to new insights into understanding brain effective connections in healthy subjects and subjects with a neurological disease. Moreover the proposed constrained model is not limited to fMRI and diffusion volumes, but it can also be applied to different domains where structural and time varying data is generated such as two-photon *Ca*^2+^ imaging (Sheikhattar et al., 2018), electroencephalography, and metabolic positron emission tomography (Hampel et al., 2011).

## Acknowledgments

We are thankful to Prof. Daniele Marinazzo for the interesting insights on how to compute Granger causality given our particular case.

## Data and code availability

All used data are publicly available online at http://fcon1000.projects.nitrc.org/.

The relevant code is publicly available at https://github.com/alecrimi/effective connectivity toolbox.

## Conflict Of Interest

The authors declare no conflict of interest.

## Authors Contribution

A.C., V.M. and D.S. designed research; A.C. and L.D performed research; A.C., F.S. and D.S. analyzed data; and A.C. and D.S. wrote the paper.

## 6. Supporting Information

**S.Figure 1:**
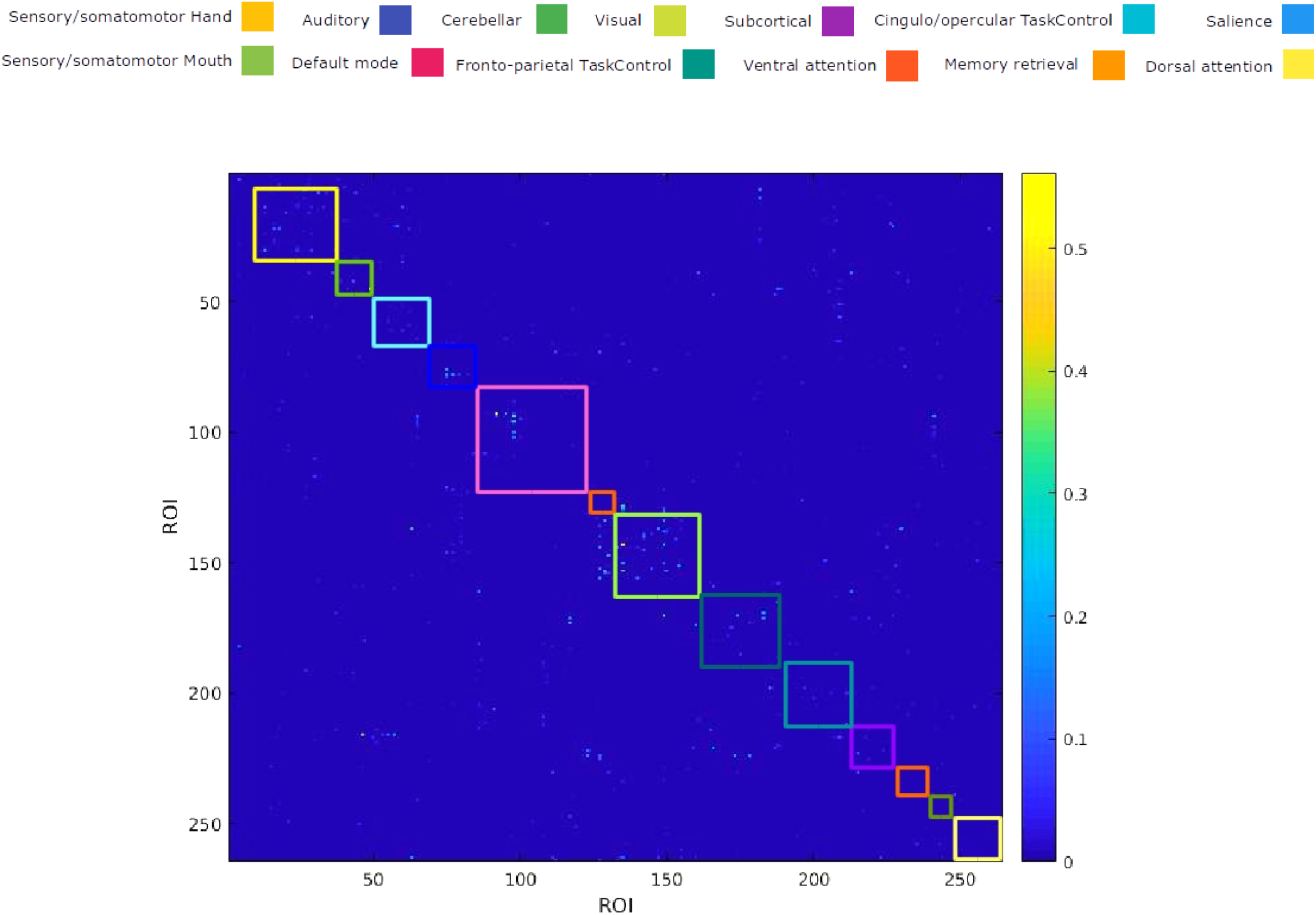
Granger causality matrix showing the ROIs with directed influence across the whole dataset according to p-values smaller than 0.05 in the F-distribution.

**S.Figure 2:**
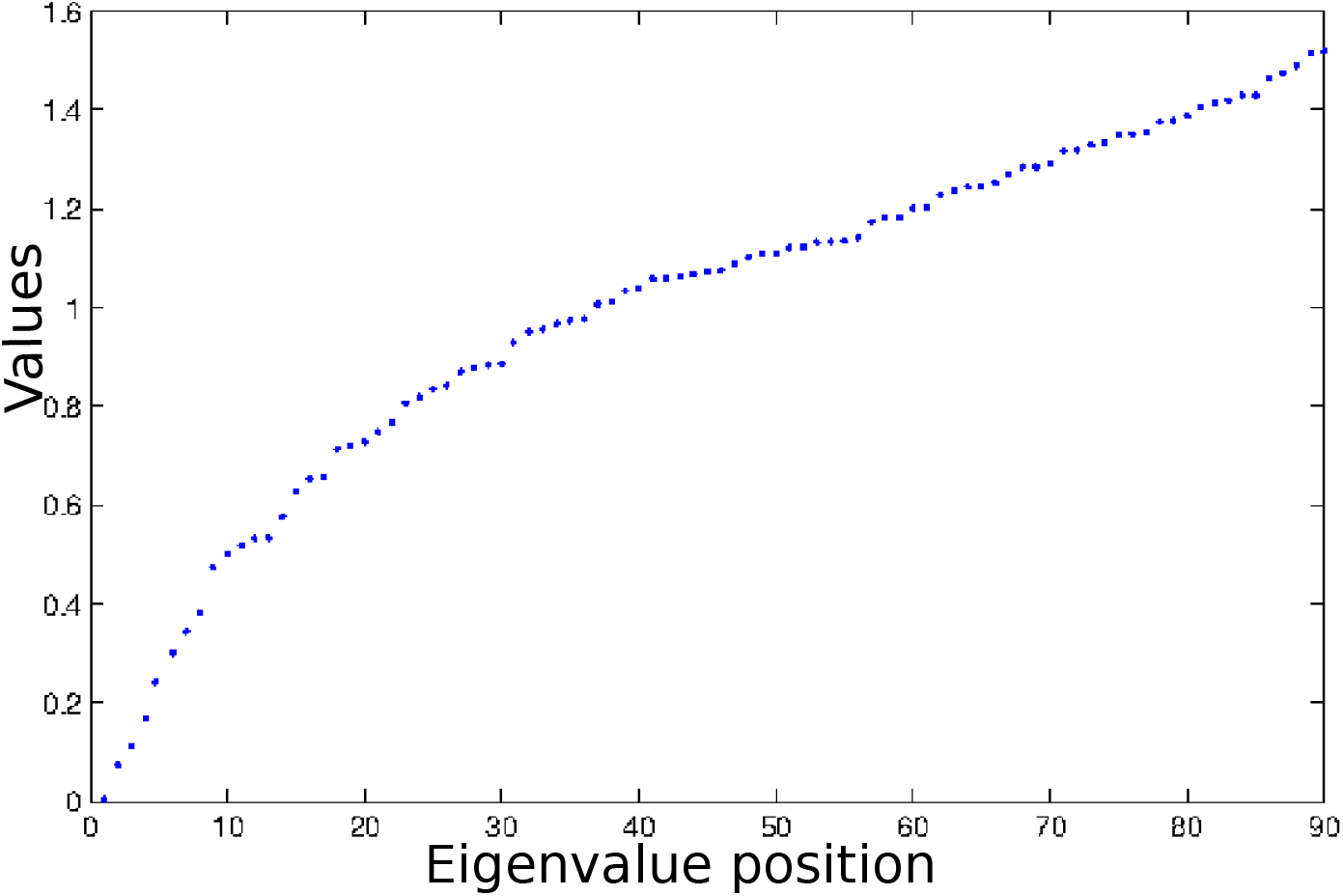
Eigengap that allows to identify the present clusters for the aforementioned experiment. It is visible that the largest gap is present at the 8th eigenvalue.

## Notes

http://www.alessandrocrimi.com/effconn/

### Competing Interest Statement

The authors have declared no competing interest.

### Summary of Updates

more experiments, change text and added 2 figures

